# ChIPSeqSpike: A R/Bioconductor package for ChIP-Seq data scaling according to spike-in control

**DOI:** 10.1101/269118

**Authors:** Nicolas Descostes, Aristotelis Tsirigos, Danny Reinberg

## Abstract

**Motivation:** Chromatin Immuno-Precipitation followed by Sequencing (ChlP-Seq) is used to determine the binding sites of any protein of interest. ChIP-Seq data suffer from being more qualitative than quantitative. The recent use of Spike-in controls along with the standard protocol tackled this problem. However, no dedicated tool is available for a robust evaluation of this new ChIP-seq approach

**Results:** We developed ChIPSeqSpike, an R/Bioconductor package that enables ChIP-Seq spike-in normalization, assessment and analysis. Ready to use scaled bigwig files and scaling factors values are obtained as output. ChIPSeqSpike also provides tools for ChIP-Seq spike-in assessment and analysis through a versatile collection of graphical functions.

**Availability:** The package is implemented in R (as of version 3.4) and is available from Bioconductor at the URL: https://www.bioconductor.org/packages/3.7/bioc/html/ChIPSeqSpike.html, where installation and usage instructions can be found.

## 1 Introduction

Chromatin Immuno-Precipitation followed by Sequencing (ChIP-Seq) is used to determine the chromatin binding sites of any protein of interest, such as transcription factors or histones with or without a specific modification, at a genome scale (Barski *et al.* 2007; Park 2009). ChIP-Seq entails: treating cells with a cross-linking reagent such as formaldehyde; isolating the chromatin and fragmenting it by sonication; immuno-precipitating with antibodies directed against the protein of interest; reversing cross-link; DNA purification and amplification before submission to sequencing. These many steps can introduce biases that make ChIP-Seq more qualitative than quantitative. Different efficiencies in nuclear extraction, DNA sonication, DNA amplification or antibody recognition can present a challenge in distinguishing between true differential binding events and technical variability.

This problem was addressed by using an external spike-in control to monitor technical biases between conditions (Orlando *et al.* 2014; Bonhoure *et al.* 2014; Chen *et al.* 2016). Exogenous DNA from a different non-closely related species is inserted during the protocol to infer scaling factors. This was shown to be especially important to reveal global histone modification differences, that are not caught by traditional downstream data normalization techniques, such as in the case of histone H3 lysine-27 trimethyl (H3K27me3) upon Ezh2 inhibitor treatment (Egan *et al.* 2016) or histone H3 lysine 79 dimethyl (H3K79me2) upon EPZ5676 inhibitor treatment (Orlando *et al.* 2014; Fig. 1).

**Fig. 1.**
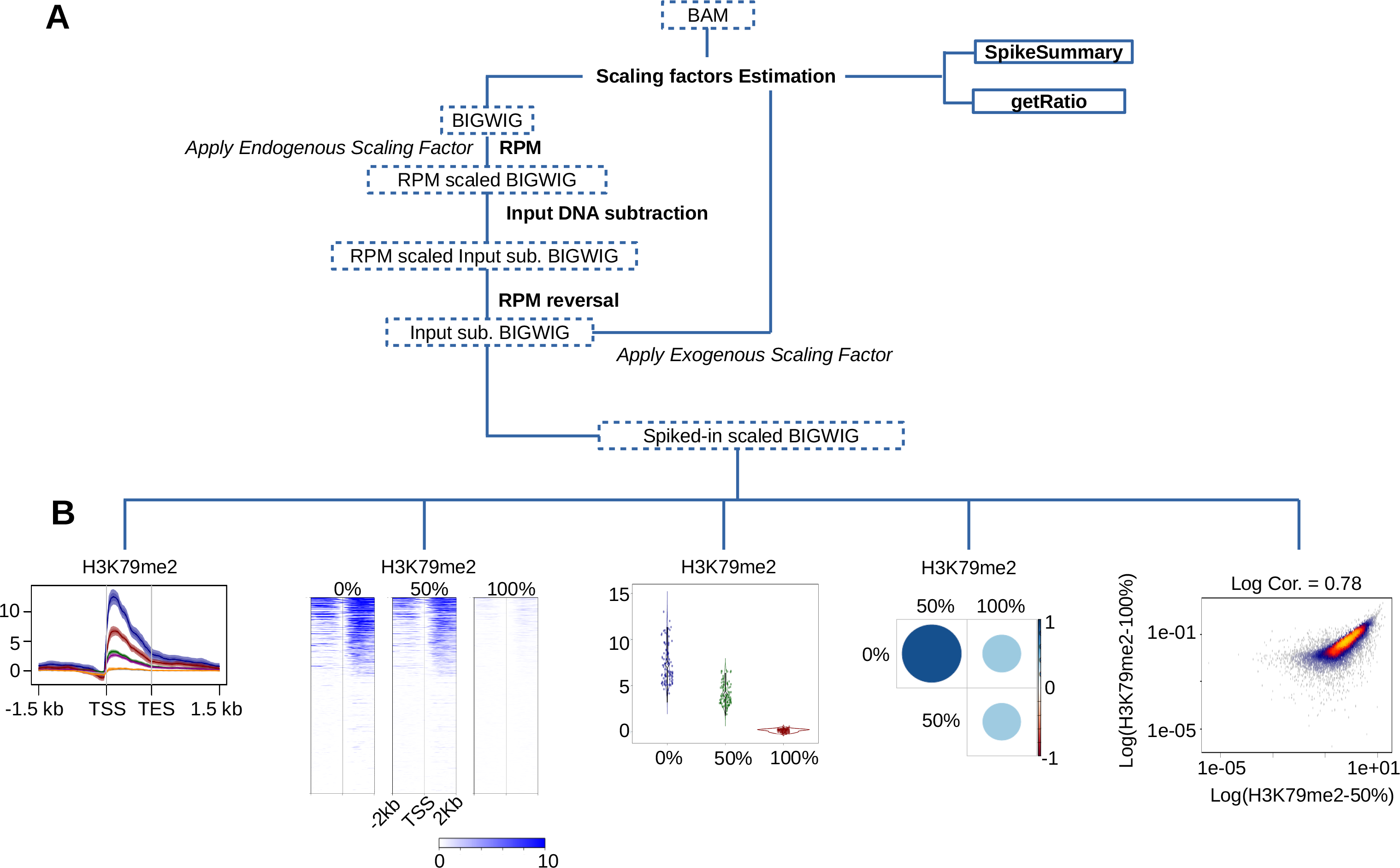
The ChIPSeqSpike package: Illustration of H3K79me2 levels upon 0%, 50%, and 100% EPZ5676 inhibitor treatment. Data references are available in supplementary information. **(A)** Diagram illustrating the different processing steps required for spike-in normalization of ChIP-Seq data. **(B)** Illustration of different graphical representations provided by ChIPSeqSpike. Here H3K79me2 ChIP-Seq data in human Jurkat cells upon treatment with different percentages of EPZ5676 inhibitor is illustrated.

Here we present ChIPSeqSpike, an R/Bioconductor package that provides tools for ChIP-Seq spike-in normalization, assessment and analysis through a versatile collection of processing and graphical functions.

## 2 Features

### 2.1 Data processing

The spike-in normalization procedure consists of 4 steps: Read Per Million (RPM) scaling, input DNA subtraction, RPM scaling reversal and exogenous spike-in DNA scaling (Fig. 1A).

The input files consists of BAM and BIGWIG files obtained after alignment on the endogenous and exogenous genome. Input DNA control files are also needed. All normalization steps can be performed independently to offer precision or at once through the usage of the “SpikePipe” wrapping function.

### 2.2 Summary and controls

The quality of the experiments can be assessed with the “SpikeSummary” and “getRatio” functions (Fig. 1A).

The first one returns the different scaling factors for each experiment as the number of reads contained in BAM files. The second one assesses the percentage of exogenous DNA relative to the endogenous DNA. This percentage should be in the 2-25% range (Orlando *et al.* 2014). ChIPSeqSpike throws a warning if this boundary is not met. In theory, having more than 25% exogenous DNA should not affect the normalization whereas having less than 2% is usually not sufficient to perform a reliable normalization.

### 2.3 Plots

ChIPSeqSpike offers visual assessment of the data with meta-profiles, heatmaps, boxplots, correlation, and heatscatter plots (Fig. 1B).

The data at a particular step of the normalization process can be integrated in a modularity manner and output into publication quality figures. Dedicated options also enable comparison of a same dataset through the different normalization steps described in section 2.1. Finally, ChIPSeqSpike offers versatile graphical options to customize boxplots and correlation plots and also provides the possibility of using different clustering techniques for heatmap representations.

### 2.4 Boost mode

Each processing step follows a read and write procedure. Reading and writing BIGWIG and BAM files can be time consuming especially when dealing with large amounts of data. ChIPSeqSpike takes t is into account by providing a “boost” mode that keeps data loaded into GRanges objects. This is particularly suitable for users having access to High Performance Computing facilities and wishing to speed up the full workflow.

## 3 Conclusion

ChIPSeqSpike is to our knowledge the first bioinformatic tool fully dedicated to ChIP-Seq spike-in data normalization. It offers a wide range of possibilities for data processing but also visualization. It offers a fast way to process the data through different modes and provides publication quality figures through a well-defined set of functions.

*Conflict of Interest:* none declared.

